# Reduced axon caliber in the associative striatum of the *Sapap3* knockout mouse

**DOI:** 10.1101/2021.01.12.426339

**Authors:** E. Lousada, M. Boudreau, J. Cohen-Adad, B. Nait Oumesmar, E. Burguière, C. Schreiweis

## Abstract

Pathological repetitive behaviors are a common feature of different neuropsychiatric disorders such as obsessive-compulsive disorder or Gilles de la Tourette syndrome. The *Sapap3* knockout mouse (*Sapap3*-KO) is the current reference model used in translational psychiatry to study co-morbid repetitive behaviors, presenting both compulsive-like as well as tic-like behaviors. Consistent with clinical and fundamental research literature relating compulsive-like symptoms to associative cortico-striatal dysfunctions and tic-like symptoms to sensorimotor cortico-striatal dysfunctions, abnormalities comprising both circuits have been described in this mouse model. Findings reported on these mice point towards not only macro-, but also micro-circuitry deficits, both of which can be affected by neuronal structural changes. As such, in the present study, we aimed to investigate structural changes in associative and sensorimotor striatal areas that could affect information conduction. We used AxonDeepSeg, an open-source software to automatically segment and measure myelin thickness and axon caliber, and found that axon caliber, the main contributor for changes in conduction speed, is specifically reduced in the associative but not the sensorimotor striatum of the *Sapap3*-KO mouse. This smaller axon caliber in *Sapap3*-KO mice is not a general neuronal feature of this region, but specific to a subpopulation of axons with large caliber. These results point to a primary structural deficit in the associative striatum, affecting signal conduction and consequent connectivity.

## Introduction

Repetitive, ritualistic actions are a core feature of animal behavior, including humans, which allow for efficient learning and automatization of behavior^1–3^. However, several neuropsychiatric and neurodevelopmental disorders, amongst them Gilles de la Tourette syndrome (TS) and obsessive-compulsive disorder (OCD) or autism spectrum disorder (ASD), are characterized by a pathological expression of such repetitive behaviors (RBs), in the form of tics, compulsions, or “stimming” behaviors, respectively.

Cortico-striatal circuits are the principal anatomical region that underlies the development of RB^4,5^. These circuits follow a parallel, mostly segregated, structural-functional organization of limbic, associative, and sensorimotor pathways of information processing^6–8^. The so-called associative circuits comprise the associative cortical areas, such as the lateral orbitofrontal cortex (lOFC), projecting to the dorsomedial striatum, the homologue of the primate caudate nucleus; while the so-called sensorimotor circuits comprise sensorimotor cortical areas, such as primary motor cortex and supplementary motor cortex, projecting to the rodent dorsolateral striatum, which corresponds to the primate putamen. Different types of RBs are thought to differently recruit each loop^9^: motor tics, the main symptom of TS patients, have been described to be more related to alterations in the sensorimotor loop^10–13^. On the other hand, RBs consisting of a more complex sequence of actions or rituals, as is the case of compulsive-like behaviors, are frequently performed by OCD patients in response to intrusive thoughts or obsessions. These cortico-striatal loops, which have been consistently described in connection with compulsive-like RBs, are referred to as the associative cortico-striatal loops^14–18^. Despite this putative anatomical segregation of different types of RBs, a great percentage of comorbidity has been reported between TS and OCD patients, which points towards the existence of a common anatomo-functional ground^19–22^.

Rodent RBs have been extensively studied in translational psychiatric approaches, with a particular focus on excessive self-grooming as a behavioral marker for compulsive behaviors^23–26^. Self-grooming is an innate rodent behavior that consists in a highly stereotyped sequence of rostro-caudal movements, essential for hygiene maintenance and overall well-being of the animal^23,24^. An aberrant manifestation of such a complex chain of movements is useful to study the mechanisms underlying the regulation and dysregulation of complex motor outputs such as compulsive-like behaviours. Currently, the reference mouse model for pathological RBs is the *Sapap3*-KO mouse^27–30^. These mice lack the synapse-associated protein 90/postsynaptic density protein 95 associated protein 3 (Sapap3), which is highly expressed in the cortex and striatum. The *Sapap3*-KO mouse model has been mainly reported and studied for its disproportionate and injurious levels of self-grooming, its increased anxiety, and its behavioral rescue with fluoxetine, a first-line treatment for OCD patients^27^. Consistently with clinical observations that the associative cortico-striatal loops are implicated in the emergence of compulsions, studies in this mouse model have shown a significant increase in the baseline firing rates of the medium spiny neurons (MSNs) in the major cortical input area of the associative circuits, the associative/centromedial striatum^28^. To further confirm the implication of the associative cortico-striatal circuits, the authors have shown that the manipulation of such circuits can influence the expression of self-grooming behaviors: optogenetic stimulation of lOFC terminals in the associative/centromedial striatum restored self-grooming levels.

More recently, a behavioral and pharmacological study of RBs in this mouse model has additionally detected tic-like RBs^31^. Tic-like behaviors correspond to simple, sudden, and rapid motor manifestations, described as head jerks and body twitches in the *Sapap3*-KO mice. Aripiprazole administration, a first-line treatment for tic-like symptoms in TS, was sufficient to drastically reduce this phenotype. In line with the hypothesis that the sensorimotor circuits are implicated in more simple motor behaviors, substantial input into the centromedial striatum coming not only from associative cortical areas such as lOFC but also from the secondary motor area (M2), rodent homologous of the primate supplementary motor cortex, has been demonstrated. The observed substantially increased input from M2 onto both MSNs and parvalbumin-positive (PV) interneurons in the centromedial striatum suggests an aberrant macrocircuitry involving, not only associative, but also sensorimotor cortico-striatal loops^30^. The implication not only of the associative but also sensorimotor cortico-striatal loops is further supported by the finding of synaptic alterations in both circuits in young adult *Sapap3*-KO mice: cingulate area 1/M2 cortical projections to the dorsomedial striatum (DMS), and primary motor/M2 cortical projections to the dorsolateral striatum (DLS)^32^. Indeed, the presence of a mixed nature of RBs and affected associative and sensorimotor circuits in the *Sapap3*-KO mice are in line with the clinical studies reporting comorbidity of tic- and compulsive-like symptoms in both OCD and TS patients^19–22^.

While numerous studies provide indications for dysfunctions on the macro-circuitry level of associative and/or sensorimotor cortico-striatal connectivity in connection with pathological RBs^27,28,30,32,33^, important dysfunctions on the micro-circuitry level within the associative striatum have equally been shown to substantially contribute to both electrophysiological as well as behavioral malfunctions. Evidence for micro-circuitry contributions comes from studies, not only on this mouse model, but also on other mouse models of RBs, as well as studies with neuropsychiatric patients. Notably, PV interneurons, which are crucially implied in regulating striatal activity through a powerful mechanism of fast-forward inhibition^34,35^, are reduced in number in the associative/centromedial striatum of the *Sapap3*-KO mouse and optogenetic elevation of PV-firing rescued striatal hyperactivity as well as aberrant behavioral response inhibition^28^. Studies on other mouse models of RBs have reported similar findings: for example, the *Cntnap2*-KO mouse exhibits lower PV expression levels within PV interneurons^36,37^. In the same line, postmortem analysis of striatal regions of TS patients has shown a reduced and altered distribution of these cells within the striatum^38,39^.

In both macro- and micro-circuitry connectivity, proper conductivity can be affected or modulated by neuronal structure and integrity. Different parameters can contribute to a deficient neuronal connectivity. On the one hand, on a cellular level, axonal structure can affect the speed of signal propagation^40^. On the other hand, cell density and ratio can affect the integration and modulation of information on a population level; for example, a reduced number of PV interneurons fails to regulate striatal activity in the context of RBs^12,28,36^. Axonal structure, in its turn, can vary according to the axon caliber and its respective myelination. Minor alterations in the caliber of the axon, or the thickness of the myelin that insulates it, provide significant changes in conduction speed by altering the resistance to the conduction of the electrical signal^40,41^. The velocity of the signal conduction is, in its turn, determinant for either strengthening or weakening the connection between two synaptically connected neurons, being thus able to modulate neuroplasticity^42,43^. Therefore, it is rather intuitive that changes in these structural parameters have been resurfaced as a potential neuroplasticity mechanism involved in learning and behavior^44–46^. Furthermore, these structural parameters have been shown to contribute to deficient neuronal signal propagation in circuits affected by major psychiatric disorders such as schizophrenia or ASD: hypomyelination specific to cortical PV interneurons has been reported in a rat model of schizophrenia^47^. Additionally, patients with ASD, a developmental behavioral disorder frequently characterized by the presence of RBs in the form of “stimming” behaviors, display a smaller axon caliber in the myelinated axons of the corpus callosum when compared to healthy controls^48^.

Different neuroimaging studies have found structural connectivity deficits in OCD patients^49,50^. However, such non-invasive methods carry the disadvantage of not being able to specify the deficit that is taking place. As far as we are informed, structural connectivity has never been assessed in a rodent model for tic- and/or compulsive-like behavior. Therefore, here, we raised the question whether structural connectivity is affected in a comorbid model of tic- and compulsive-like RBs, using the current reference mouse model of these type of RBs, the *Sapap3*-KO mouse model,. Hereby, we covered both candidate regions that have previously been characterized as affected both in human patients with tic- and/or compulsive-like behaviours as well as according animal models: the associative (AS) and the sensorimotor striatum (SMS), which form part of the associative or sensorimotor cortico-striatal circuits, respectively. In both candidate regions, we morphologically characterized myelinated axons in the *Sapap3*-KO mouse model, measuring both axon caliber and myelin thickness as the determinant parameters of axon conductivity. To do so, we took advantage of the open-source software AxonDeepSeg^51^, which automatically segments axon and myelin compartments using deep learning methods and performs various morphometric measurements (axon caliber, myelin thickness, g-ratio). AxonDeepSeg is fast (a few seconds per image), allowing the analysis of hundreds of images in only a few minutes. As myelin coverage is done by oligodendroglial cells, i.e. the glial cells that wrap around the fibers to produce the myelin sheaths, we additionally screened this cell type in the same regions of interest for potential abnormalities. Finally, we analyzed the density of myelinated axons, a proxy for the density of myelinated cells, in *Sapap3*-KO mice given previous reports on cell count changes in the dorsomedial striatum of these mice^28^.

## Results

### Axon caliber is diminished in the associative but not in the sensorimotor striatum of *Sapap3*-KO mice

In order to assess striatal structural integrity of the *Sapap3*-KO mouse model of tic- and compulsive-like behaviors, we characterized axon area and myelin thickness, two parameters that crucially contribute to axonal conductivity (Fig. 1A). Hereby, we distinguished between associative and sensorimotor striatal regions, i.e. striatal areas receiving projections from the respective cortical areas^52^, which are located at the medial or lateral striatal boundary along the dorso-central continuum, respectively (Fig. 1B). Taking advantage of a deep learning algorithm, AxonDeepSeg, which is capable of automatically, rapidly and accurately segmenting and measuring myelinated axons from electron microscopy data in an unbiased manner^51^ (Fig. 1C), we assessed the properties of approximately 150 axons per region and animal. Applying such tool, we measured axon area, and myelin thickness, i.e. the difference between outer and inner myelin layer (Fig. 1C); the latter was extracted via the according outer and inner axonal surfaces. Additionally, we calculated the g-ratio, a classical measurement that describes the myelin thickness in proportion to the caliber of the axon by dividing the radius of the inner layer by the radius of the outer layer (n = 4 per genotype; AS: median_WT_ = 0.80 vs. median_KO_ = 0.79; Mann Whitney U: U = 12, p-value = 0.34; SMS: median_WT_ = 0.81 vs. median_KO_ = 0.80; Mann Whitney U: U = 13, p-value = 0.20; Fig. S1, upper panels).

**Figure 1:**
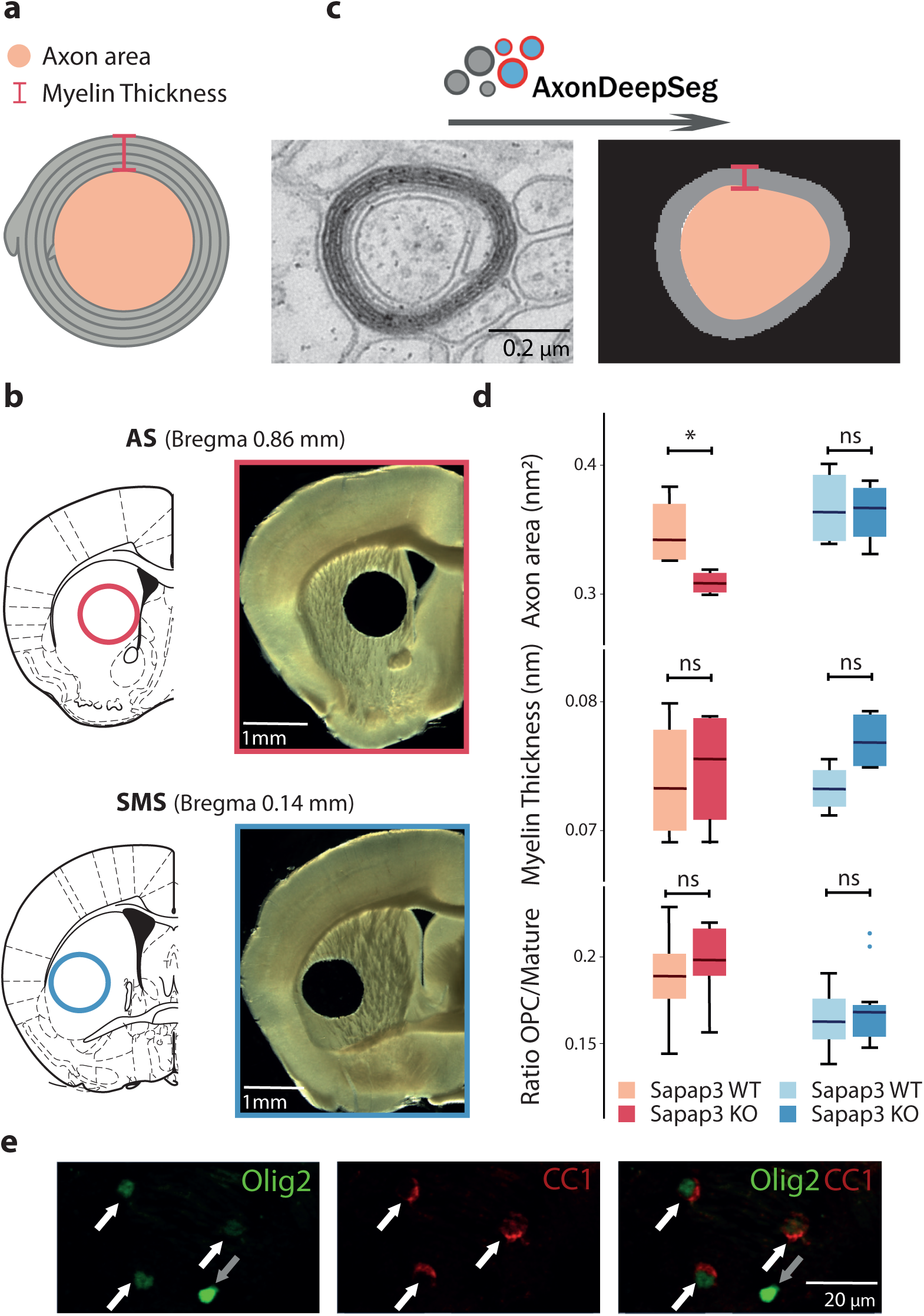
Axon caliber, and not myelination, is altered in the associative striatum of the *Sapap3*-KO mouse. **a**. Scheme of a myelinated axon highlighting the parameters of interest, the axon area (light pink) and myelin thickness (grey, delimitations in dark pink). **b**. Schemes (left panels) and stereomicroscopic images (right panels) of coronal brain slices at target bregma levels illustrating punch-extraction of the regions of interest of the associative (red outlines) and the sensorimotor striatum (blue outlines). **c**. Schematic illustration of the proceeding by the AxonDeepSeg algorithm. Electron microscope images (100x acquisition) containing myelinated axons (left panel) are automatically detected and measurements of axon surface and myelin thickness extracted (right panel). **d**. Axon area, myelin thickness, and ratio between immature and mature oligodendroglial cells in Sapap3-KO (darker colors) and wild-type controls (lighter colors) in the associative (red shades) and the sensorimotor striatum (blue shades). Box-whisker plots illustrate 25^th^ and 75^th^ percentiles respectively, and medians. *: p < 0.05; ns = non-significant. **e**. Exemplary split channel (left and middle panel) and merged channel (right panel) immunofluorescence image (x20 magnification) of immature (Olig2+/CC1-; grey arrows) and mature (Olig2+/CC1+; white arrows) oligodendrocytes. Olig2 signal is pseudocoloured in green, CC1 signal in red. AS = associative striatum; SMS = sensorimotor striatum.

We found that axon area was significantly diminished in the associative striatum of the *Sapap3*-KO mice in comparison with their wild-type age matched controls (median_WT_ = 0.34 vs. median_KO_ = 0.31; Mann Whitney U: U = 16, p-value = 0.03). However, there was no difference in axon area in the sensorimotor striatum (median_WT_ = 0.36 vs. median_KO_ = 0.37; Mann Whitney U: U = 9, p-value = 0.89) (Fig. 1D, upper panels). On the other hand, myelin thickness in *Sapap3*-KO mice was unaltered in either striatal region (AS: median_WT_ = 0.07 vs. median_KO_ = 0.08; Mann Whitney U: U = 8, p-value = 1; SMS: median_WT_ = 0.07 vs. median_KO_ = 0.08; Mann Whitney U: U = 2 p-value = 0.11) (Fig. 1D, center panels). These results suggested that myelination per se might not be affected in *Sapap3*-KO mice. However, in order to utterly exclude potential changes on myelination itself, we furthermore histologically analyzed the cells from the oligodendroglial lineage, i.e. those cells, which wrap around and directly myelinate the fibers. Thus, we performed immunostainings in order to quantify the overall existing population of oligodendroglial cells, using oligodendrocyte transcription factor 2 (Olig2) as a cell marker; as well as only the mature pool of oligodendrocytes, i.e. those oligodendrocyte forms, which actively perform myelination. For the latter we applied the criterion of a positive double-immunolabeling for both Olig2 as well as adenomatous polyposis coli clone CC1 (CC1) (Fig. 1E, white arrows). Thus, cells that are only Olig2-positive corresponded to immature oligodendroglial cells (Fig. 1E, gray arrows). In line with the results of unaltered myelin thickness in associative and sensorimotor striatal regions of *Sapap3*-KO mice, as obtained via the AxonDeepSeg approach, no significant differences were observed neither in the number of immature (n = 12 per genotype; AS: median_WT_ = 91.56 vs. median_KO_ = 94.38; Mann Whitney U: U value = 60, p-value = 0.51; SMS: median_WT_ = 75.90 vs. median_KO_ = 71.95; Mann Whitney U: U = 76.5, p-value = 0.82; Fig. S1 center panels) nor in the number of mature, i.e. myelinating oligodendrocytes (n = 12 per genotype; AS: median_WT_ = 517.18 vs. median_KO_ = 497.72; Mann Whitney U: U = 80, p-value = 0.67; SMS: median_WT_ = 441.85 vs. median_KO_ = 445.24; Mann Whitney U: U = 77, p-value = 0.80; Fig. S1 bottom panels). We additionally assessed the ratio of immature versus mature oligodendrocytes, as a different proportion would give insight into eventual maturation deficits of this cell line and also did not detect significant differences (n = 12 per genotype; AS: median_WT_ = 0.19 vs. median_KO_ = 0.20; Mann Whitney U: = 115, p-value = 0.16; SMS: median_WT_ = 0.16 vs. median_KO_ = 0.17; Mann Whitney U: U = 131, p-value = 0.62) (Fig.1D, bottom panels).

The striatum is the main input region of cortical projections and *Sapap3* deletion has shown to specifically affect cortico-striatal projections^33^. Hence, in order to assess whether the detected difference in axon caliber is specific to the associative striatum or whether overall cortico-striatal circuitry is affected, we expanded the analysis of axon caliber and myelin thickness to the lateral OFC (lOFC) as well as to the primary and secondary motor cortex (M1/M2), i.e. those cortical regions that provide respective major inputs into the associative and sensorimotor striatum (Fig. S2A). We found no significant difference in neither axonal parameters in lOFC as well as M1/M2 (n = 4 per genotype; Axon area – lOFC: median_WT_ = 0.46 vs. median_KO_ = 0.46; Mann Whitney U: U = 8, p-value = 1; M1/M2: median_WT_ = 0.55 vs. median_KO_ = 0.50; Mann Whitney U: U = 13, p-value = 0.20; Fig. S2B upper panels; Myelin thickness – lOFC: median_WT_ = 0.08 vs. median_KO_ = 0.08; Mann Whitney U: U = 5, p-value = 0.49; M1/M2: median_WT_ = 0.08 vs. median_KO_ = 0.08; Mann Whitney U: U = 7, p-value = 0.89; Fig. S2B center panels). In the same line, we did not find any differences in the ratio of oligodendroglial cells (n = 12 per genotype; lOFC: median_WT_ = 0.27 vs. median_KO_ = 0.30; Mann Whitney U: U = 63, p-value = 0.62; M1/M2: median_WT_ = 0.31 vs. median_KO_ = 0.30; Mann Whitney U: U = 72, p-value = 1; Fig. S2B bottom panels).

### The reduction in axon caliber arises from a subpopulation of axons

Having detected a significant decrease in the axon caliber of the *Sapap3*-KO mice in the associative striatum, we pondered whether this reduction would arise from an overall reduced caliber of myelinated axons, or rather from a reduction specific to a particular range of axon populations. As a third option, we furthermore considered the possibility of a reduction in the number of a specific subpopulation of axons characterized by a large caliber, hereby skewing the overall caliber size to a smaller average in the *Sapap3*-KO mice. In order to investigate these three different possibilities, we plotted the probability distribution of all axon areas, segregating *Sapap3*-KO and *Sapap3*-WT groups, using the Raincloud plot algorithm^53^. The obtained probability density plot suggested a bimodal distribution in the axon area of *Sapap3*-WT mice, a distribution that appeared blunted in the *Sapap3*-KO (Fig. 2A). Attending to the considerable heterogeneity of striatal cells and the observed bimodal distribution, we fitted a two-component Gaussian mixture model to our data in order to test for the potential presence of two different clusters. Indeed, this model confirmed the visual impression of a bimodal distribution and detected the presence of two clusters: the first, larger cluster contained 85%of axons of the *Sapap3*-KO group, and 82% of the *Sapap3*-WT group (Cluster 1), and the other cluster contained 15% and 18% of axons (Cluster 2), respectively (Fig. 2B). We verified between-subject comparability of the distributional profiles (Fig. S3). Axon areas in Cluster 1, which comprises the vast majority of the axons, were similar between *Sapap3*-KO and *Sapap3*-WT mice (n_WT_ = 514, n_KO_ = 508; median_WT_ = 0.26 vs. median_KO_ = 0.24; Mann Whitney U: U = 131604.5, p-value = 0.29). However axons of *Sapap3*-KO mice assigned to Cluster 2 were significantly smaller than those of their wild-type age matched controls (n_WT_ = 113, n_KO_ = 90; median_WT_ = 0.67 vs. median_KO_ = 0.58; Mann Whitney U: U = 7986.0, p-value = 6,52 x 10^−4^) (Fig. 2B). These results thus suggest that the observed reduction in axon caliber in the associative striatum of *Sapap3*-KO mice is specific to a small subpopulation of axons characterized by larger axons, here assigned to Cluster 2; reduced axon caliber is thus not a generalized feature of all myelinated axons in the associative striatum.

**Figure 2:**
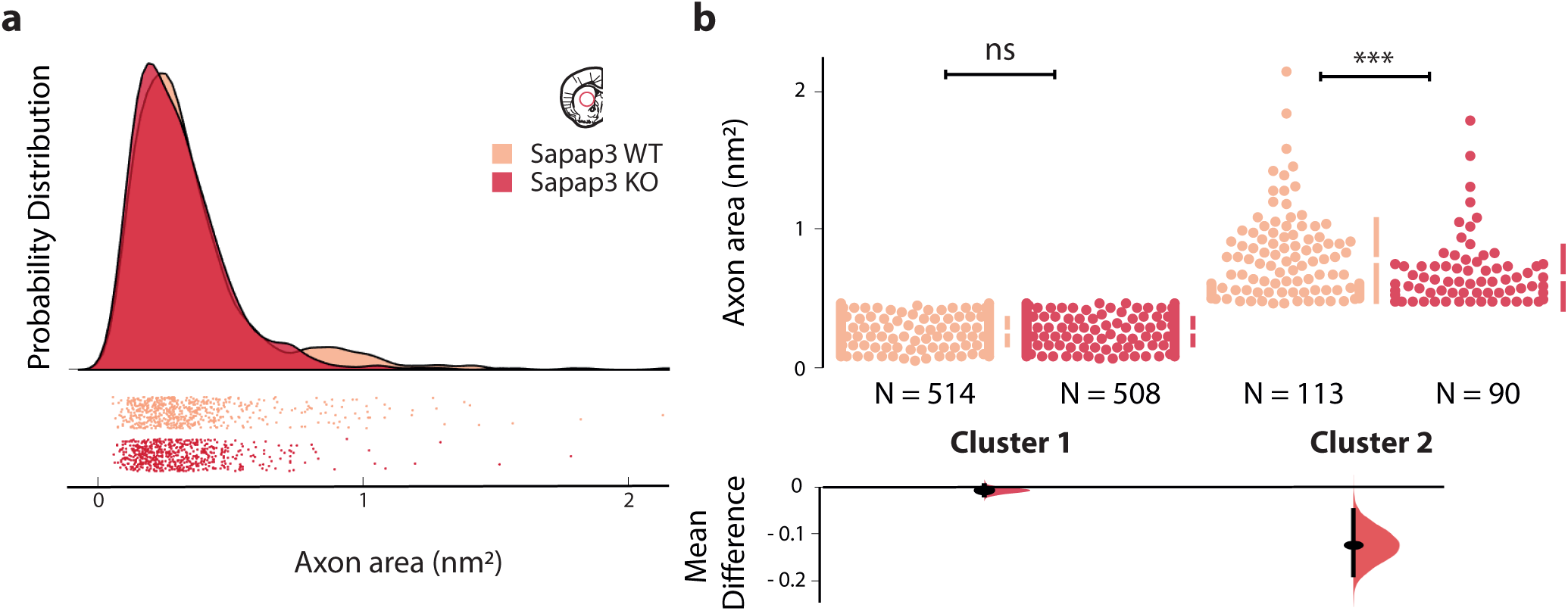
Axon area difference in the associative striatum of *Sapap3*-KO mice originates from a subpopulation of neurons. **a**. Individual axon area data points of the associative striatum of *Sapap3*-KO and wild-type mice (below) and raincloud plot of their probability distribution (on top). Separate peaks in the probability distribution suggest the presence of two clusters in both *Sapap3*-KO and wild-type mice. **b**. Myelinated axons (individual data points; upper panel) form two clusters (cluster 1, left; cluster 2, right) according to their axon area in the associative striatum of *Sapap3*-KO and wild-type mice (color coding as above) as confirmed by fitting a two-component Gaussian mixture model (upper panel). The axon area of myelinated axons in cluster 2 is significantly reduced in *Sapap3*-KO compared to wild-type mice (lower panel). *Sapap3*-KO mice are depicted in dark, wild-type mice in light red. ***: p < 0.001; ns = non-significant.

However, a third hypothesis remained to be explored: was the genotype difference in Cluster 2 due to an overall reduction of axon caliber within that cluster? Or could this difference be explained by the relative lack of a subpopulation of large axons within Cluster 2, hereby skewing axon caliber averages within that cluster to a smaller average in *Sapap3*-KO mice? The latter possibility would thus be reflected in a decreased overall number of myelinated axons in the associative striatum of *Sapap3*-KO mice. Hence, we quantified and averaged the density of myelinated axons in 50 transmission electron microscopy images per animal in the associative striatum, comparing again *Sapap3*-KO and wild-type age matched controls. Attending to the fact that the striatum is traversed by innumerous fibers of the internal capsule, which largely project outside of the striatum, we visually excluded the bundles of the internal capsule (Fig. 3A). We did not detect any difference in the density of myelinated axons in the associative striatum between *Sapap3*-KO and *Sapap3*-WT animals (n = 4 per genotype; median_WT_ = 20.80 vs; median_KO_ = 21.41; Mann Whitney U: U = 6, p-value = 0.69; Fig. 3B). Hence, this suggests that a lack of a subgroup of axons with large caliber cannot account for the axon caliber difference in the associative striatum of *Sapap3*-KO mice. In other words, the observed genotype-dependent difference in axon caliber in the associative striatum is likely due to an overall reduction in axon caliber of Cluster 2.

**Figure 3:**
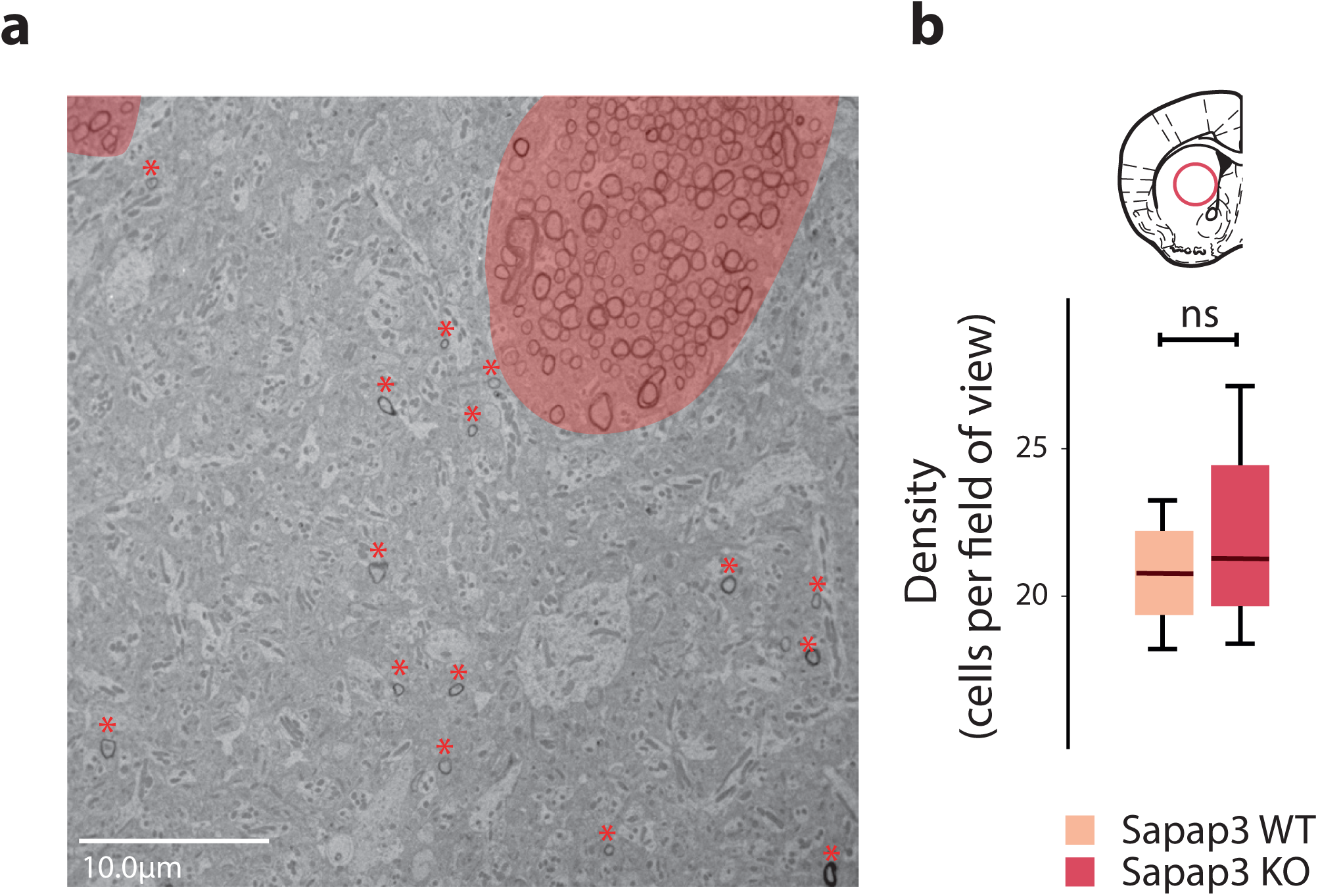
Density of myelinated axons is not altered in the associative striatum of *Sapap3*-KO mice. **a**. Exemplary electron microscopy image (x6.2 magnification), highlighting the internal capsule, which was excluded from analysis (light pink), and quantified axons beyond the internal capsule (dark pink asterisks) **b**. Density of myelinated axons in the associative striatum of *Sapap3*-KO (dark red) and wild-type mice (light red). Box-Whisker plots illustrate the 25^th^ and 75^th^ percentile and the median. ns = non-significant.

## Discussion

In the present study, we demonstrated that structural connectivity is affected in the *Sapap3*-KO mice, a mouse model of comorbid tic- and compulsive-like behaviors^31^. This deviation consists in a reduced axon caliber of myelinated fibers specifically in the associative striatum; such a structural deficit was not detected neither in the sensorimotor striatum nor in upstream cortical regions, namely lOFC and M1/M2, respectively. Furthermore, we provide evidence that this reduction in axon caliber affects only a small cluster of axons characterized by a large caliber, representing less than 20% of the overall number of myelinated axons in the associative striatum. Myelination itself, however, as assessed through myelin thickness and histological quantification of the myelin-generating oligodendrocytes, seems unaffected. Even though myelin thickness is dependent on the diameter of the axon^54^, it has been shown that diameter does not regulate myelination alone^55^, thus being able to be independently altered.

Axon caliber is directly correlated with conduction speed. In fact, it has been shown that fiber caliber, by increasing axial resistance^56^, is the main contributor to variations in the conduction speed^57^. In its turn, a diminished conduction speed affects synaptic transmission and thus transsynaptic communication with downstream cells^42,43^. Indeed, it has been demonstrated in primate motor callosal neurons that smaller axon diameter corresponds to smaller synaptic bouton size and consequent smaller neurotransmission release^58^. Hence, the reduced axon caliber in the associative striatum of *Sapap3-*KO mice further corroborates a connectivity deficit present in this region as previously suggested by other studies using in vitro electrophysiology^27,30,32,33^.

The *Sapap3*-KO mouse presents a behaviorally complex phenotype composed of a spectrum of different types of RBs^31^. On the one hand, this mouse model has been firstly characterized by its disproportionate and disrupted sequences of self-grooming, its increased anxiety, and its responsiveness to fluoxetine^27^. As such, and considering the fact that abnormal self-grooming, due to its phasic, chained, and syntactic nature, has been used to understand hierarchical motor outputs, the *Sapap3-*KO mouse has been considered a model of exclusively compulsive-like behaviors. In addition, consistently with a large part of the clinical literature on OCD^14,15,17,18^, abnormalities in the cortico-striatal associative circuits such as described for patients with OCD have also been described in this mouse model, with particular focus on the corticostriatal pathway between the lateral OCF and the associative striatum^59,60^ However, a large part of the clinical literature pronounces the heterogeneity of OCD and an important comorbidity between tic-like and compulsive-like behaviors in both TS and OCD patients. Recently, *Sapap3*-KO mice have been found to additionally present tic-like movements such as simple head twitches and body jerks, behaviors, which are rescued with aripiprazole administration^31^; these findings are consistent with yet another part of the clinical literature describing high comorbidity between OCD- and TS-like symptoms^19–22^. As such, in a mouse model that spans such a complex spectrum of tic- and compulsive-like RBs, we were surprised to find that axon caliber was affected only in the associative striatum, while the sensorimotor striatum was found unaltered. How could altered axon caliber specific to the associative striatum explain such a co-morbid behavioral phenotype, for which, in the most straightforward manner, both an associative as well as a sensorimotor circuitry component could be expected?^10–12,14–18^

One possible explanation for the presence of tic-like movements in this mouse model could be due to the strong sensorimotor cortical input from M2 into the associative striatum in addition to associative cortical areas ^30^. Indeed, as Corbit et al. described, this input is strengthened in the *Sapap3-*KO mice, suggesting a maladaptive reinforcement from sensorimotor relative to associative areas that would lead to aberrantly elevated motor output. Given aberrantly elevated input from M2 onto the associative striatum as shown by Corbit et al., a concomitant decrease in associative circuits, as suggested in our study, could further corroborate an imbalance between sensorimotor and associative circuits, which has been suggested previously for patients and animal models of RBs ^61,62^. Such an imbalance could for example facilitate the disruption of the processing of complex and sequential motor outputs, in favor of breaking them into simpler, shorter, and purely motor behaviors.

Currently, aberrant connectivity in the associative striatum, here observed as reduced axon caliber can be explained in two ways^27,30,32,33^. A first explanation is based on dysfunctional macrocircuitry of cortico-striatal loops. Concretely, in this scenario, the observed reduction in axon caliber in a small cluster of striatal axons might arise from a population of cortico-striatal projection neurons. Given that axonal caliber is positively related to synaptic strength^42,43^, the here observed structural phenotype of a decrease of axon caliber in the associative striatum is in line with and corroborates a previously reported decrease in cortico-striatal synaptic strength in the associative striatum^30,32^. No such structural alterations were observed in cortical regions providing major input into the associative striatum, including lOFC and M2^30,32,52^. Cortic-striatal tracing studies will be useful in the future to further investigate this structural macro-circuitry hypothesis.

Alternatively, the observed structural deviations in the associative striatum could arise from alterations in the striatal micro-circuitry, namely a diminished feed forward inhibitory network of striatal PV interneurons. A reduced axonal caliber arising specifically from the subpopulation of PV interneurons would be consistent with previously observed diminished cortico-striatal synaptic activity in general^58^, and with decreased numbers of PV interneurons specifically^28,32^. Altered striatal micro-circuitry in form of a reduced inhibitory PV interneuronal network has been described both in patients with aberrant repetitive behaviors^38,39,63^ as well as in the striatum of several mouse models^36,37,64^ including the associative striatum of Sapap3-KO mice^28,32^. PV interneurons, even though representing a small percentage of the total number of striatal cells, form a strong inhibitory blanket across the striatum. Alterations in these cells, either by reduction in size, or through a deficient morphology and connectivity with downstream cells, reduces their feedforward inhibitory power, failing therefore to finely orchestrate the MSNs in their role of behavioral control^28,65^. Alternatively, a very similar idea has been rising that it is not the number of PV interneurons that is altered in case of aberrant repetitive behaviors, but expressions levels of parvalbumin. Indeed, neuronal PV expression levels are altered given its modulatory role in synaptic plasticity^66^. Hence, an observed decrease in the number of PV interneurons might be a consequence of thresholding PV expression levels in immunohistochemistry cell counts^67,68^. Reduced concentration of this protein has been confirmed in several mouse models with aberrant repetitive behaviours^37,69^. Taken together, there are several indications that the here detected smaller axon caliber in a subpopulation of associative striatal neurons in Sapap3-KO mice might arise from deviations in the pool of striatal PV interneurons. However, surely, there is not direct translation of the observed structural phenotype to a reduced number of PV interneurons for several reasons. First, in our study we additionally measured the density of myelinated axons in the associative striatum without detecting a genotype-dependent difference. This indicates that there might be no change in the number of neuronal cells with myelinated axons.

Second, we found the smaller axon caliber to be specific to a subpopulation of myelinated axons making up about 15-18% of the entire population of myelinated axons in the region. The population of PV interneurons, however, corresponds to only about 1% of the total number of striatal neurons^70^. Having said this, it needs to be noted that, as far as we are informed, this proportion has never been assessed considering only striatal cells with myelinated axons; for example dopaminergic striatal afferents have reported to not be myelinated^71,72^. Furthermore, the axonal arborization of striatal PV interneurons is extremely dense, thus making it possible to measure the same axonal tree of a single even though rare PV interneuron several times within one section^73^.

And, third, a portion of striatal cholinergic (ChAT) interneurons has also been reported to be myelinated^74^. As PV interneurons, these ChAT interneurons, even though sparse, highly contribute to the modulation of MSNs^38,39,75^ and could potentially be deficient in our mouse model. Indeed, ChAT interneurons have been described to be reduced in number in TS patients^38^, and their partial ablation produces tic-like behaviours in mice^12^, as well as compulsive social behaviors^76^.

In summary, our study corroborates previous findings of deficits within the associative striatum of the *Sapap3*-KO mouse, here assessed for the first time using a structural, deep learning-based approach. This study thus contributes a novel and complementary aspect to the discussion of deviations in macro- and microcircuitry in the context of RBs in clinical as well as fundamental studies.

## Materials and methods

### Animals

Adult *Sapap3-*KO and wild-type age matched controls (*Sapap3-*WT) (n = 32; aged 5-11 months) of C57BL/6J background, were generated in heterozygous breeding trios and housed at the animal facilities of the Paris Brain Institute in Tecniplast ventilated polycarbonate cages under positive pressure with hard-wood bedding and provided with ad libitum food and water. The temperature was maintained at 21–23 °C and the relative humidity at 55 ± 10% with a 12-h light/dark cycle (lights on/off at 8am and 8pm, respectively). All experiments were approved by the French ministry of research under the agreement number (APAFIS) #1418-2015120217347265. Founders for the *Sapap3*-KO colony were kindly provided by Drs. Ann M Graybiel and G. Feng, MIT, Cambridge, USA. Genotyping was performed as in the original publication^27^.

### Selection of regions of Interest

In order to determine the delimitations of the associative (AS) and sensorimotor striatum (SMS), we analyzed anterograde projection patterns documented for the lateral orbitofrontal cortex (lOFC) and primary and secondary motor cortices (M1/M2) in the Allen Mouse Brain Connectivity Atlas^52^. Bregma levels resulting from this hodological screening of the two striatal regions of interest were thus defined at +0.86 for the associative striatum (AS) and at +0.14 for the sensorimotor striatum (SMS)^77^. Bregma levels of the lOFC and M1/M2 areas corresponded to bregma = +2.68 and bregma = + 1.94, respectively.

### Electron Microscopy

Animals (n = 4 per genotype) were deeply anesthetized with pentobarbital (200mg/kg) and transcardially perfused with fresh 5% glutaraldehyde (GA)/5mM CaCl_2_. Brains were collected, post-fixed overnight in 5% GA at 4°, then rinsed and kept overnight in 0.12M phosphate buffered solution (PB) (pH=7.4).

Brains were cut in 200µm slices on the vibratome; slices comprising the regions of interest were selected and their left-hemispheric part punched under a stereomicroscope using a biopsy punch with plunger of 1mm in diameter (PFM Medical). Punched samples were incubated with 4% osmium tetroxide in 100mL sodium cacodylate buffer for one hour at RT and then carefully rinsed with sodium cacodylate buffer, three times. Next, samples were immersed in 5% uranyl acetate in 100mL sodium cacodylate buffer for one hour and carefully rinsed in sodium cacodylate buffer, three times. The punches were then dehydrated in increasing concentrations of ethanol (50%, 70%, 90%, and twice 100%) for five minutes, and finally twice in 100% acetone for ten minutes. In order to prepare the samples for epoxy resin embedding, they were first immersed overnight, at 4°C, in a solution of 50:50 acetone and epoxy resin, then transferred into embedding molds and immersed in pure resin for two hours, and finally left to polymerize at 60°C for 48hours in a new dose of epoxy resin. Samples were then cut into 70nm slices on an ultramicrotome, placed onto a Transmission Electron Microscope (TEM) grid and immersed in lead citrate for fifteen minutes.

Samples were imaged at x100.0k for quantifications of axon caliber, myelin thickness and g-ratio, and at x6.2k for the quantification of the density of myelinated axons in the Transmission Electron Microscope (HITACHI 120kV HT7700, camera AMT XR41-B). In order to be specific on the quantification of local striatal cells, myelinated axons within the fiber bundles that cross the striatum, i.e. axons corresponding to the internal capsule, were excluded from analysis.

### Electron Microscopy Analysis

#### Myelin thickness and axon area/caliber measurements

We quantified approximately 150 myelinated axons per region and animal using the AxonDeepSeg^51^. In order to train the algorithm, we used a subset of TEM images and manually drew the inner and outer myelin layers using Fiji, ImageJ2. We then created an 8-bit image delineating the axon caliber area (i.e. area inside of the myelin inner layer) in white, the myelin area (i.e. area between the inner and the outer myelin layers) in grey, and background area (i.e. area outside of the outer myelin layer) in black. These ground truth masks were then provided to the algorithm together with the TEM images to train the model. After training, TEM images were fed into the trained model for automatic segmentation. At this step, we visually checked and corrected eventual artifacts, before running the complementary algorithm that measured the area of the detected and segmented regions as inferred by pixel size. To validate the model, we manually analyzed all AS images, confirming results as obtained through the algorithm.

#### Quantification of myelinated axons

For each animal, we analyzed 50 images per region of interest (AS, SMS, lOFC, M1/M2) and manually assessed the number of myelinated axons in each image, using Fiji, ImageJ2^78,79^.

In order to be specific on the quantification of local striatal cells, myelinated axons that form part of the internal capsule, i.e. fiber bundles that cross the striatum, were excluded (Fig. 3**a**).

### Immunohistochemistry

Animals (n = 12 per genotype) were deeply anesthetized with pentobarbital (200mg/kg) and transcardially perfused with 2% PFA in 1xPBS. Fixed brains were dissected and post-fixed overnight in 2% PFA at 4°C. Samples were immersed in 15% sucrose in 1xPBS for 24 hours, and subsequently immersed in 30% sucrose in 1xPBS for 48 hours. Samples were then embedded in Tissue-Tek O.C.T. compound and frozen on dry ice. Brains were subsequently sliced coronally in 12µm sections on a cryostat, transferred on a coated glass slide (Superfrost Ultra Plus, Thermo Scientific), and stored at − 80°C.

For immunohistochemical labeling, we started by heat-inducing antigen retrieval with unmasking solution and then, after cooling down at RT for twenty minutes, we rinsed the samples with 1X PBS three times for five minutes. Samples were then blocked in 4% BSA/0.1% Triton X-100 for 1 hour.

The primary antibodies against Oligodendrocyte transcription factor 2 (Olig2) mouse IgG2a, nuclear marker for all cells of the oligodendroglial lineage, and against adenomatous polyposis coli clone CC1 (CC1) mouse IgG2b, a cytosolic marker specific for mature oligodendrocytes, were diluted in 4% BSA/0.1% Triton X-100 with a concentration of 1/500 and 1/100 respectively, and incubated overnight at 4°C. Samples were rinsed with 1X PBS and incubated with the secondary antibodies rat anti-Mouse IgG2a 488 and goat anti-Mouse IgG2b 555 Alexa in a 1/2000 dilution, for 1 hour at RT, in the dark. The slides were rinsed under agitation in the dark, mounted in Mowiol mounting medium, coverslipped, and left overnight to dry at RT, protected from light.

Fluorescent samples were imaged in a Zeiss Axio Observer 7 at x20 magnification. The ROIs we chosen visually by an experienced user and corresponded to one field of view (665,60µm x 665,60µm). We imaged both right and left hemispheres of two slices per region and animal, separated by 72µm, summing up to a total of four images per region for each mouse.

### Immunohistochemistry Analysis

For the quantification of oligodendrocyte density, we used Fiji, ImageJ2^78,79^ and manually counted all cells that were positively stained for Olig2 and CC1. Positive immunolabeling for both Olig2 and CC1 (Olig2+/CC1+ cells) allows for the identification of mature oligodendrocytes, while positive immunolabeling of Olig2, and negative labeling of CC1 (Olig2+/CC1-cells) allows for the identification of immature cells of the oligodendrogial lineage.

## Data Analysis and Statistics

All data and statistical analysis were performed using Matlab R2017b. Given small sample size, we performed exclusively non-parametric Mann-Whitney-U tests. We furthermore used raincloud plots^53^ as well as estimation plots^80^ for visualization of our data, two open-source software packages available on GitHub. Cluster analysis was done by fitting a Gaussian mixture model with two components (2 clusters) to our data^81^.

## Data Availability

The datasets and analyses of this study are available from the corresponding author on reasonable request.

---

**Supplementary Figure S1:**
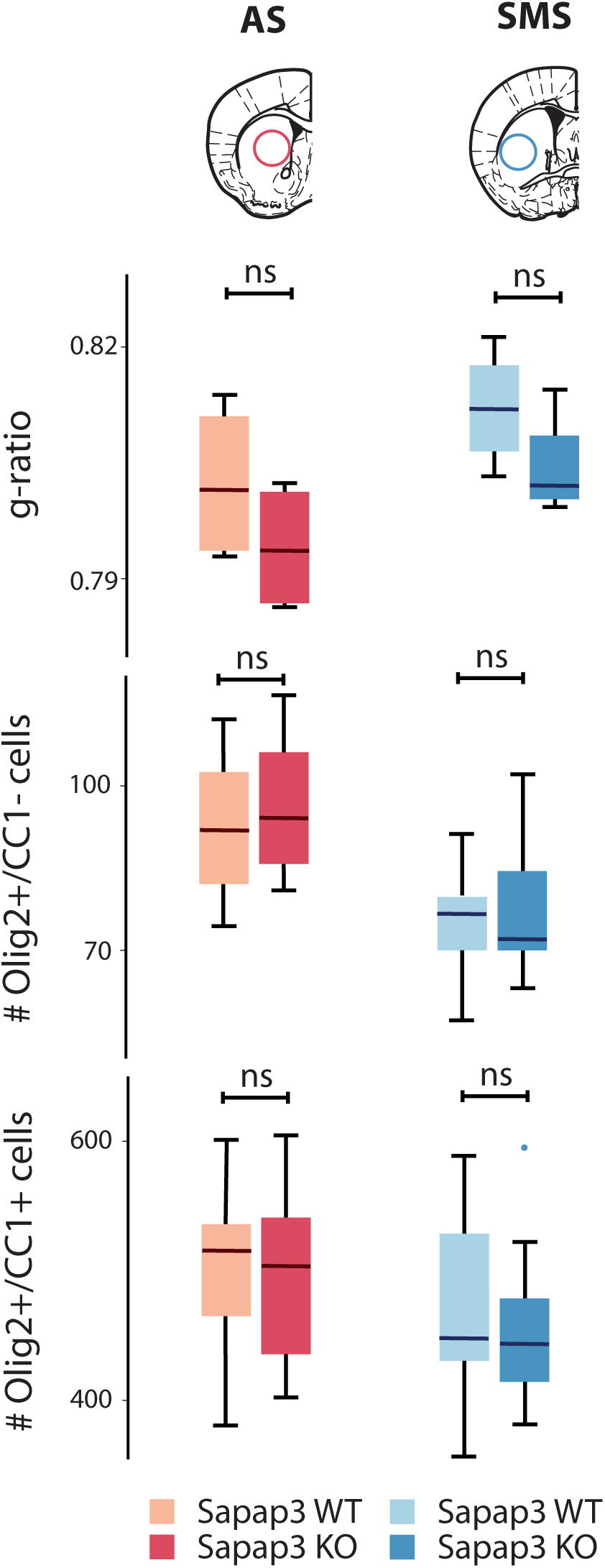
G-ratio and oligodendroglial cells are not altered in the striatum of the *Sapap3*-KO mouse. G-ratio (upper panel), number of Olig2+/CC1-, i.e. immature oligodendroglial cells (center panel), and number of Olig2+/CC1+ cells, i.e. mature oligodendrocytes (bottom panel) in *Sapap3*-KO (darker colors) and wild-type controls (lighter colors) in the associative (red shades) and the sensorimotor striatum (blue shades). Box-Whisker plots illustrate 25^th^ and 75^th^ percentiles respectively, and medians. ns = non-significant

**Supplementary Figure S2:**
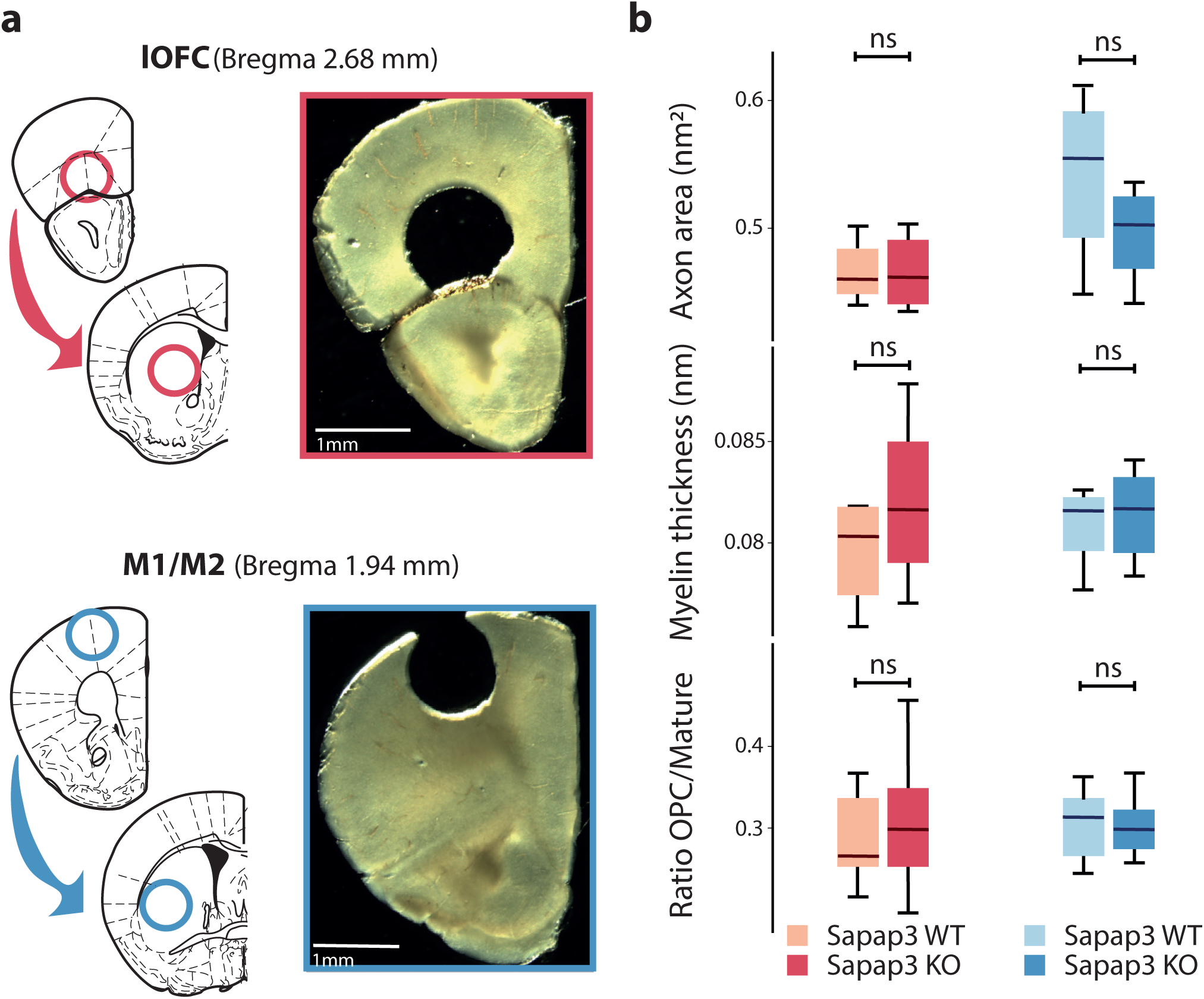
Axon area, myelin thickness, and oligodendroglial cells are not altered in the cortical input regions of the Sapap3-KO mouse. **a**. Schemes (left panels) and stereomicroscopic images (right panels) of coronal brain slices at target bregma levels illustrating punch-extraction of the regions of interest, which provide important input to the associative or sensorimotor striatum, i.e. lateral OFC (red outlines) and the M1/M2, respectively (blue outlines). **b**. Axon area, myelin thickness, and ratio between immature and mature oligodendroglial cells (upper, center, and bottom panels, respectively) in *Sapap3*-KO (darker colors) and wild-type controls (lighter colors) in the OFC (red shades) and the M1/M2 (blue shades). Box-Whisker plots illustrate 25^th^ and 75^th^ percentiles respectively, and medians. ns = non-significant.

**Supplementary Figure S3:**
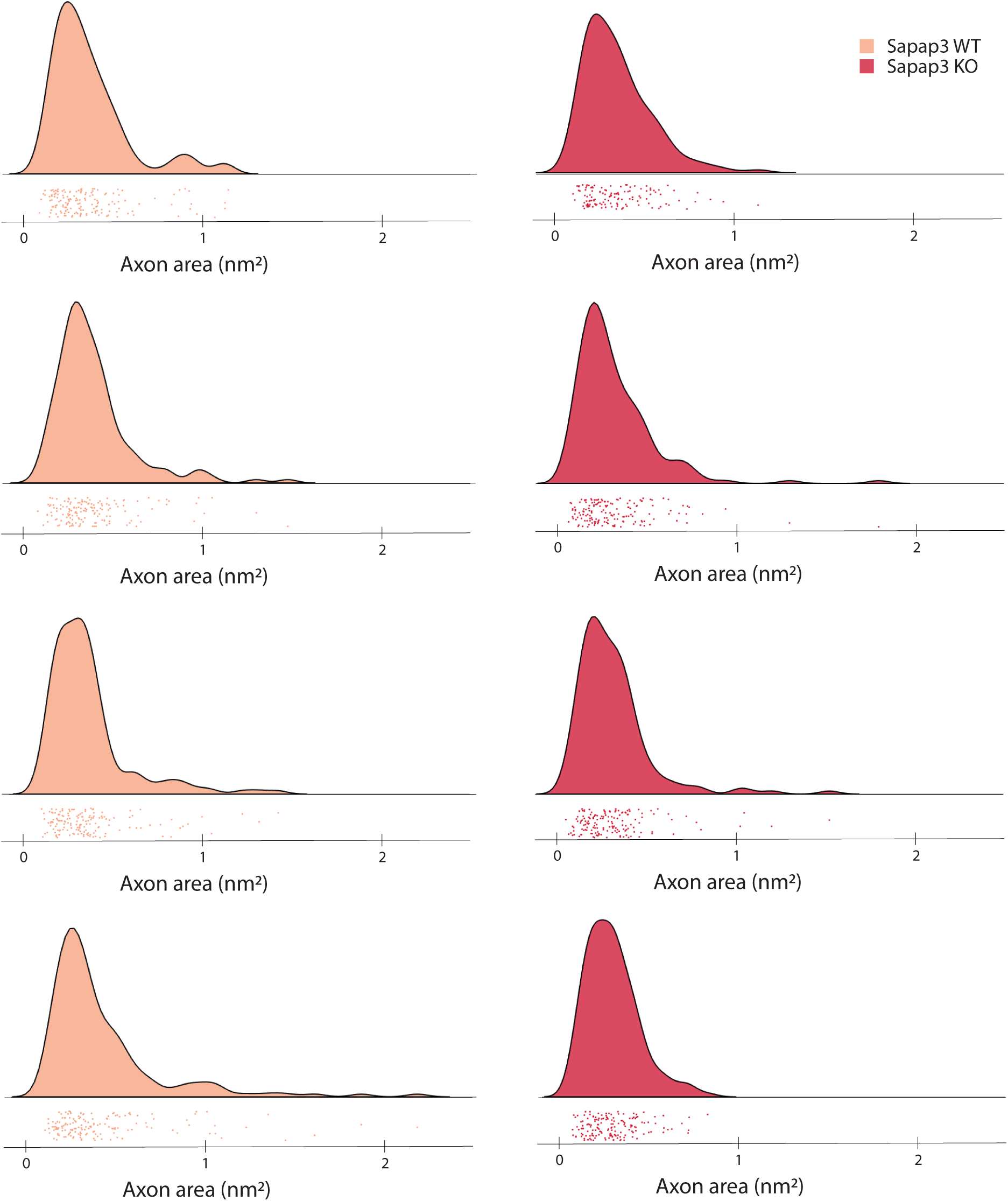
Individual subject probability distribution of axon area. Individual axon area data points of the associative striatum of *Sapap3*-KO (dark pink) and wild-type mice (light pink) (below each individual plot) and raincloud plot of their probability distribution (top of each individual plot).

## Acknowledgements

All animal work was conducted at the PHENO-ICMice facility. This work furthermore benefitted from the equipment and services from the iGenSeq core facility at the ICM for the genotyping of the animals. All electron microscopy imaging was performed using the equipment of the ICM.QUANT core facility, with particular regards to all the advice and help provided by Dominique Langui. All fluorescent microscopy was conducted using the equipment at the CELIS core facility. Moreover, sample preparation and immunohistochemistry were possible using the equipment at the HISTOMICS core facility. We thank Drs. G. Feng and Ann M. Graybiel for kindly providing founders of the *Sapap3*-KO colony, Christian Perone for helping us apply AxonDeepSeg, Dr. Dorien Maas for her guidance with immunohistochemistry protocols, Sami Beaumont for all the advice on analytical and statistical tools, as well as Marine Euvrard for her help with animal experiments. This work was realized with the following funding: Agence Nationale de la Recherche (ANR-16-INSERM-SINREP, ANR-19-ICM-DOPALOOPS) (EB), the ICM Big Brain Theory Program (BBT-ACTIMYEL; EB, BN-O), the L’Oréal-UNESCO Fellowship 2016 (CS), the Canada Research Chair in Quantitative Magnetic Resonance Imaging [950-230815], the Canadian Institute of Health Research [CIHR FDN-143263], the Canada Foundation for Innovation [32454, 34824], the Fonds de Recherche du Québec - Santé [28826], the Natural Sciences and Engineering Research Council of Canada [RGPIN-2019-07244], the Canada First Research Excellence Fund (IVADO and TransMedTech) and the Quebec BioImaging Network [5886, 35450]. The core facilities were supported by “Investissements d’avenir” (ANR-10-IAIHU-06 and ANR-11-INBS-0011-NeurATRIS) and “Fondation pour la Recherche Médicale”.

## Author Contributions

<u>EL</u>: Performed research, analyzed data, and wrote the paper

<u>MB and JC-A</u>: Contributed analytic tools, and edited the paper

<u>BN-O</u>: Designed Research, contributed reagents, edited the paper

<u>E.B</u>: Designed Research, and wrote the Paper

<u>C.S</u>: Performed Research, designed Research, and wrote the Paper

